# Brain temperature affects quantitative features of hippocampal sharp wave ripples

**DOI:** 10.1101/2022.01.21.477285

**Authors:** Peter C. Petersen, Mihály Vöröslakos, György Buzsáki

## Abstract

Biochemical mechanisms are temperature-dependent. Brain temperature shows wide variations across brain states, and such changes may explain quantitative changes in network oscillations. Here we report on the coupling of various hippocampal sharp wave ripple features to brain temperature. Ripple frequency, occurrence rate and duration correlated with temperature dynamics. By focal manipulation of the brain temperature in the hippocampal CA1 region, we show that ripple frequency can be increased and decreased by local heating and cooling, respectively. Changes of other parameters, such as the rate of SPW-R and ripple duration were not consistently affected. Our findings suggest that brain temperature in the CA1 region plays a leading role in affecting the ripple frequency, whereas other parameters of SPW-Rs may be determined by mechanisms upstream from the CA1 region. These findings illustrate that physiological variations of brain temperature exert important effects on hippocampal circuit operations.

## Introduction

Hippocampal-neocortical communication is thought to be essential for different forms of cognitive behaviors, most notable for memory, imagination and planning (Maguire and Hassabis, 2011; Schacter et al., 2007). A key physiological pattern in this communication during ‘offline’ brain states (Vanderwolf, 1969) is the sharp wave-ripple complex (SPW-R; Buzsáki et al., 1992). In recent decades, SPW-Rs were the subject of extensive investigation and implicated in a plethora of functions including memory consolidation (Girardeau et al., 2009), sleep homeostasis (Miyawaki and Diba, 2016), synaptic plasticity (Norimoto et al., 2018) and metabolic regulation (Tingley et al., 2021). A particularly interesting debate revolves around the similar versus distinct features of SPW-Rs that occur in the waking resting animal and nonREM sleep (Roumis and Frank, 2015). An attractive feature of SPW-Rs is their time-compressed spike sequences of past waking experience and their potential to affect future behavior (Diba and Buzsáki, 2007; Nádasdy et al., 1999; Pfeiffer and Foster, 2013; Wilson and McNaughton, 1994). While both forward and reverse replays are present during both waking and nonREM sleep, wake and sleep SPW-Rs may have potentially different functions, given that they are embedded in different constellations of network states. In the waking animal, SPW-Rs may serve memory retrieval (Roumis and Frank, 2015), memory maintenance (Gillespie et al., 2021), stabilization of place cells (Roux et al., 2017), planning of actions and travel paths (Dupret et al., 2010; Pfeiffer and Foster, 2013), or a combination of these functions (Joo and Frank, 2018; Ólafsdóttir et al., 2018). In contrast, SPW-Rs during nonREM sleep may be critical for consolidation of long-term memories (Buzsáki, 1989), homeostatic maintenance (Miyawaki and Diba, 2016) and affecting endocrine functions (Tingley et al., 2021).

The macroscopic features of SPW-Rs, such as their incidence, amplitude, duration, and frequency, may also be different in the sleeping and waking animals. The extracellular sharp wave (SPW) is produced by synchronous transmembrane currents in the apical dendrites of CA1 pyramidal cells, which is triggered by the synchronous CA3 inputs targeting the mid str. radiatum (Buzsáki et al., 1983). The CA3 volley also excites CA1 interneurons and their interaction induces a short-lived fast oscillation (the “ripple”; 110-160 Hz) detected in the local field potential (O’Keefe and Nadel, 1978; Buzsáki et al., 1992; Stark et al., 2014; Ylinen et al., 1995). The ripple frequency is determined mainly by the local interaction of perisomatic interneurons (Stark et al., 2014) and the synchrony of pyramidal neuron spikes in each ripple wave within the CA1 region (Schomburg et al., 2012) but it is not known why ripple frequencies in the waking and sleeping brain are different. One potential explanation is brain temperature. During physiological conditions brain temperature fluctuations range 2.5°C in humans (Dijk and Czeisler, 1995), and 3°C in rodents and the strongest determinant of brain temperature is wake-sleep state rather than circadian phase (Franks and Wisden, 2021). Previous research has already established that neuronal sequences in birds, slow oscillations and sleep spindles in mammals decelerate when temperature decreases (Andersen and Moser, 1995; Csernai et al., 2019; Deboer, 1998; Franken et al., 1992; Hubbard et al., 2020; Long and Fee, 2008; Moser et al., 1993; O’Leary and Marder, 2016; Petersen and Buzsáki, 2020; Sheroziya and Timofeev, 2015) and that brain temperature regulation may have restorative functions to the brain (McGinty and Szymusiak, 1990; Franks and Wisden, 2021). To test the hypothesis that ripple features can be explained by temperature, we first examined the correlation between fluctuation of brain temperature and various parameters of SPW-Rs. To offer more direct evidence for the importance of brain temperature in regulating ripples, we artificially cooled and warmed local volume of tissue in the dorsal CA1 region. We observed a strong correlation between hippocampal temperature changes and various aspects of SPW-Rs in the sleep-wake cycle and show that local cooling and warming affect ripples.

## Results

### Hippocampal temperature correlates with brain states

Brain temperature showed wide fluctuations (~3°C) across natural behaviors (Fig. 1A-C). Waking during exercise had the highest mean temperature (36.1±0.42 °C, mean±SD), whereas the lowest temperature level was observed during sleep (nonREM 35.5±0.44 °C; REM 35.4±0.42 °C, mean±SD). The wake-sleep state variation was described quantitatively by the autocorrelogram of the temperature, corresponding to approximately ~ a 90 min cycle (Fig. 1D). The fastest brain temperature changes occurred at the transitions between brain states. The fastest change occurred at the onset of REM sleep (Fig. 1E), with an average temperature increases of >0.3°C within 2 minutes. After nonREM onset, brain temperature decreased more gradually by > 0.1°C within a few minutes (Fig. 1F), whereas wake onset showed a gradual increase (Fig. 1G). The short transient microarousals of nonREM sleep (Watson et al., 2016) did not bring about an obvious change in temperature (Fig. 1H). These brain state changes were associated with characteristic changes of the theta-delta ratio (theta range: 5 Hz - 12 Hz, delta range: 0 Hz - 4Hz) and movement (Fig. 1E-H).

**Fig. 1:**
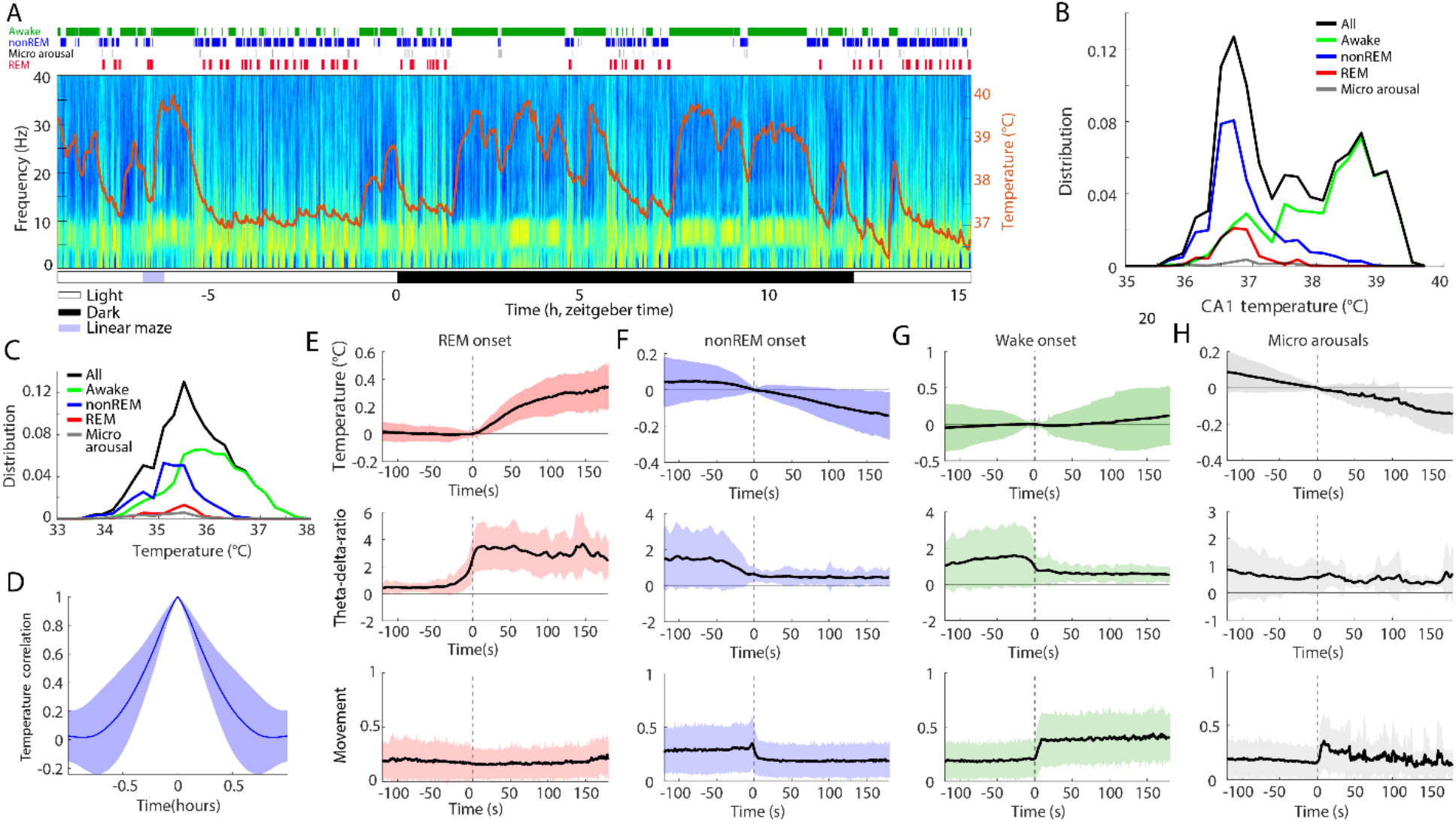
Hippocampal temperature varies across brain states. 24-hour recording of hippocampal activity and brain temperature in a chronically implanted rat. **A)** Time-power analysis of hippocampal local field potentials (LFPs) and brain temperature (orange line overlayed on spectrogram). Local field potential from the CA1 region of the hippocampus was used to calculate the time-resolved fast Fourier transform-based power spectrum. Brain state classification (Watson et al., 2016) is shown above the spectrum (awake, nonREM, micro arousals and REM, green, blue, black, and red lines, respectively). Note steep temperature rise during maze running (purple rectangle, ~ 2.5 h). **B)** Brain temperature varied ~3-4 °C within 24 hours (black), with differing distributions across brain states (awake, nonREM, micro arousals and REM, green, blue, black, and red lines, respectively). **C)** Distribution of brain temperatures across 18 recording sessions in 8 animals. **D)** Temperature autocorrelogram capturing the timescale of the temperature fluctuations. **E)** REM onset-triggered brain temperature changes (top panel), theta-delta-ratio and movement (lower panel; measured with an accelerometer on the animals’ head). **F-H)** Same panels as in E for nonREM onset (F), Wake onset (G), and Micro arousals (H).

### Hippocampal ripple metrics correlate with brain temperature in freely moving rats

Next, we characterized various hippocampal ripple metrics with the continuous brain temperature readings. Ripple frequency and the rate of SPW-R events showed large variations both across wake-sleep states as well as within nonREM and awake (Fig. 2A-F). Within long sleep episodes, a positive relationship between temperature, ripple frequency and rate of SPW-R occurrence was often visible. We observed a positive correlation between the ripple peak frequency and brain temperature (R_nonREM_ = 0.66 and R_Awake_ = 0.55; n =15 sessions). A negative correlation was observed between the ripple duration and the brain temperature (R_nonREM_ = −0.55 and R_Awake_ = −0.22; n = 15 sessions). The ripple occurrence rate was correlated with brain temperature yet with larger variance between sessions (R_nonREM_ = 0.24 and R_Awake_ = −0.11; n = 15 sessions). Quantification of these relationships showed a significant difference between nonREM and waking for ripple frequency and rate of SPW-Rs but not for their duration (Fig. 2G-I). We also calculate the temperature correlations of these same measures, which remained unchanged between nonREM and wake brain states.

**Fig. 2:**
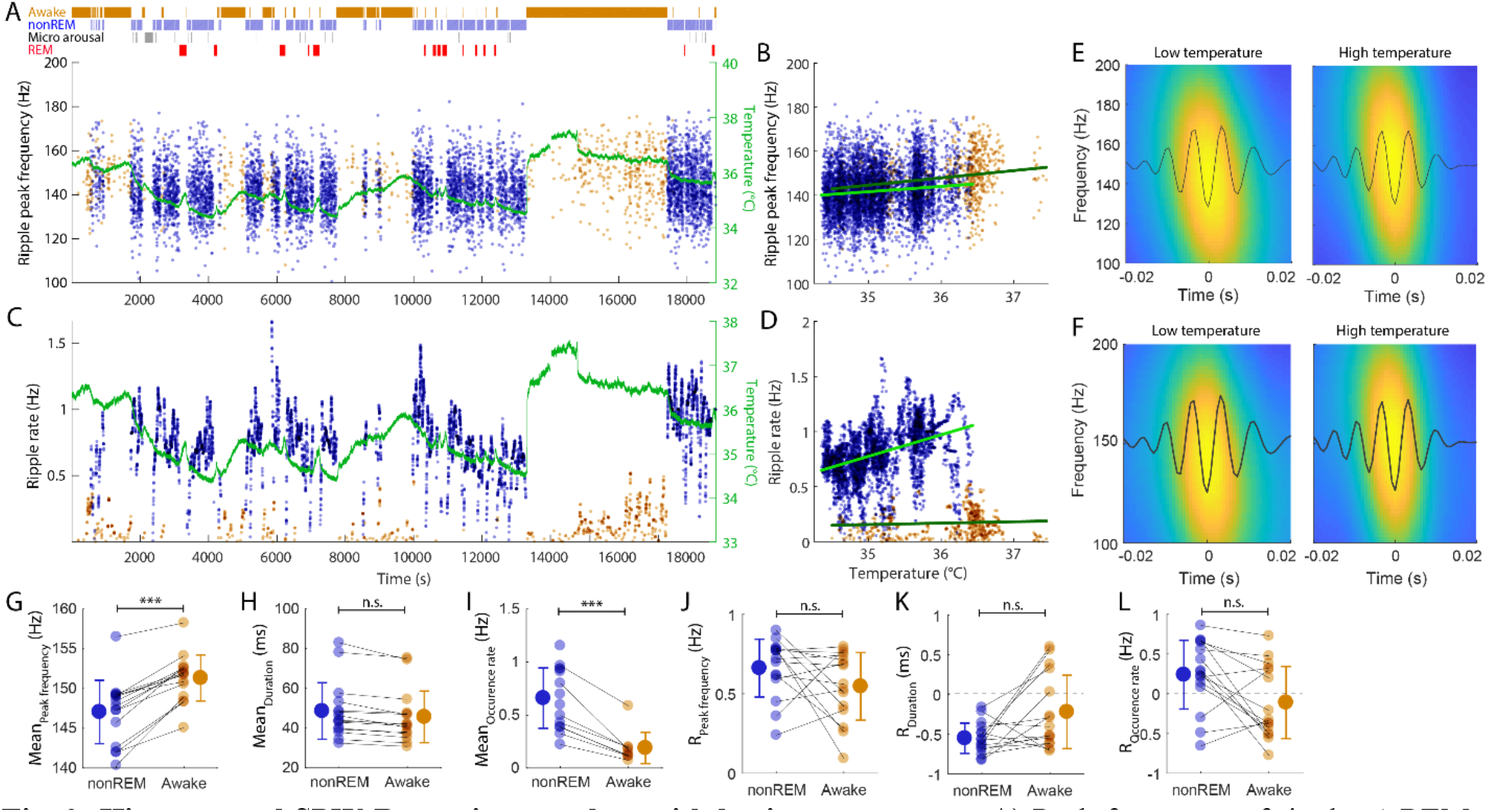
Hippocampal SPW-R metrics correlate with brain temperature. **A)** Peak frequency of ripples (nREM = blue dots; waking = brown dots) and brain temperature (green line) across time. **B)** Correlation between temperature fluctuation and peak frequency of ripples during nonREM and waking (nonREM R = 0.11/0.45, p < 0.001, slope = 2.38 Hz/°C; wake R = 0.19/0.66, p < 0.001, slope = 3.31 Hz/°C). **C)** Rate of SPW-Rs and brain temperature across time (ripple rate is calculated in 30 s intervals within brain states). Temperature line is superimposed to facilitate comparison (as in A). Note parallel change of SPW-R rate with temperature. Also note small but reliable temperature increase during REM episodes. **D)** Correlation between temperature fluctuation and the rate of SPW-R occurrence during nREM and waking (nonREM R = 0.41, p < 0.001, slope = 0.20 Hz/°C; wake R = 0.06, n.s., slope = 0.01 Hz/°C). **E)** Average ripple waveforms and wavelet maps for low (200 ripples) and high (200 ripples) temperature epochs from the session shown A-D. **F**) Average ripple waveforms and wavelet maps for low (200 ripples) and high (200 ripples) temperature epochs for all sessions. **G-I)** Peak ripple frequency, mean duration and mean rate of SPW-R occurrence during nonREM and waking. Pairs of recordings from the same session are connected. **J-L)** Correlation values between brain temperature and peak ripple frequency, duration and occurrence rate of SPW-R during nonREM and waking.

### Multivariable linear regression model prediction of ripple frequency

To assess the potential contribution of other factors, besides temperature, in describing the ripple frequency, we applied a multi variable linear regression model, taking brain temperature, the power spectrum slope, brain states (Awake, nonREM, REM and micro arousals), SPW-R rate and the theta-delta-ratio, which are changing on a similar timescale. Using a leave-one-out approach, in which we compared the performance of a linear regression model based on all parameters versus a model in which one of the parameters were left out, we found that the brain temperature was the strongest contributor, significantly higher than the other variables as quantified by the root mean squared error (RMSE; Figure 3A).

**Fig. 3:**
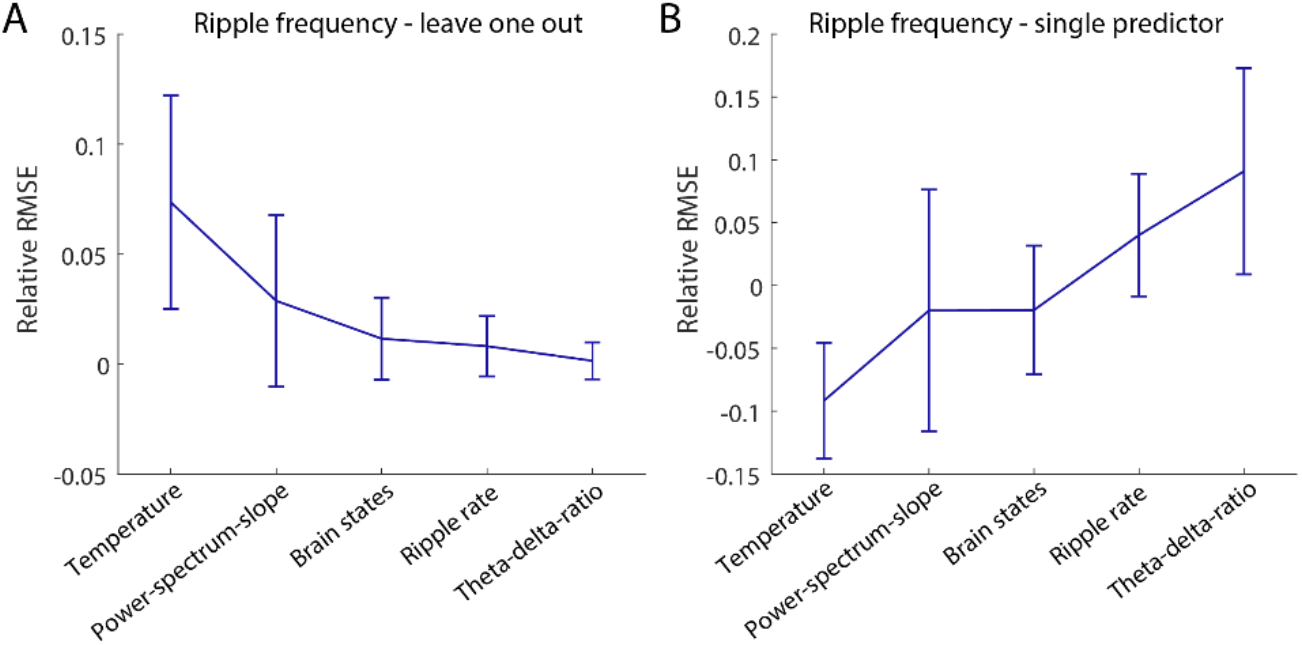
The brain temperature is the best predictor of ripple frequency. **A)** Leave-one-out prediction of ripple frequency dynamics. Root mean squared error (RMSE) difference for each held-out predictor. The held-out predictor is labelled along the X-axis. **B)** Same analysis as in (**A**) but using only one predictor at the time, showing the lowest error when predicting the ripple frequency via brain temperature (ANOTA paired test p<0.005 for all pairs). Predictors: Brain temperature, Power-spectrum-slope, brain states (awake, nonREM or REM), ripple rate, theta-delta-ratio.

To validate that the other measures were not co-variating, thus masking a potential hidden contribution to the regression model when leaving out a predictor, we also applied the same analysis using single variables. Again, the best single predictor was the brain temperature, with the lowest RMSE (Figure 3B).

### Local temperature manipulation of the hippocampus

In a further attempt to disentangle the hypothesized temperature effect on ripples from potential hidden factors, we varied the local temperature in the hippocampus. We built a device that allowed for focal cooling and heating in freely moving rats (Aronov and Fee, 2011; Long and Fee, 2008; Petersen and Buzsáki, 2020). The temperature manipulation probe consists of a silver wire (260 μm in diameter), air isolation, and a polyimide tube (Fig. 4A). The back end of the silver wire was coiled at the cold plate of a Peltier device (1.2 mm by 1.9 mm), with an added passive copper heatsink. Cooling and heating were achieved by directing current through the Peltier device. A small temperature sensor (thermistor; 280 μm in diameter) was attached at the tip of the cooling probe for continuous monitoring of the local brain temperature. The cooling probe was implanted in the CA1 region together with a silicon probe 1 mm apart (Fig. 4A-D). By reversing the current, we were able to induce both local heating and cooling within the same recording session (Fig. 4E-G). Local temperature manipulation did not affect the sleep-wave cycle or within-sleep state changes (Fig. S1-2). Local cooling (Δ_temp_ = −2.8°C) significantly lowered the ripple frequency on the ipsilateral side by approximately 1.7 Hz (n = 12 sessions, P=0.034, Wilcoxon signed rank test), whereas heating (Δ_temp_ = 3.7°C) increased ripple frequency by 1.5 Hz (Fig. 4 H; n = 12 sessions, P=0.012, Wilcoxon signed rank test). Ripple duration and the occurrence of ripple rates was not consistently affected by the local manipulation of the temperature (Fig. 4I, J). While heating increased ripple duration, cooling had no effect (Fig. 4I; n = 12 cooling sessions, P = 0.016; n = 12 heating sessions; P =0.077). In contrast, cooling had no effect on SPW-R rate, whereas heating had a small, although significant increase (Fig. 4J; n = 12 cooling sessions, P = 0.30; n= 12 heating sessions; P =0.027). Identical temperature manipulation of the contralateral hippocampus was without an effect (Fig. S3; n = 5 cooling sessions, P = 0.99, n = 8 heating sessions, P = 0.52, Two-sample Kolmogorov-Smirnov test).

**Fig. 4:**
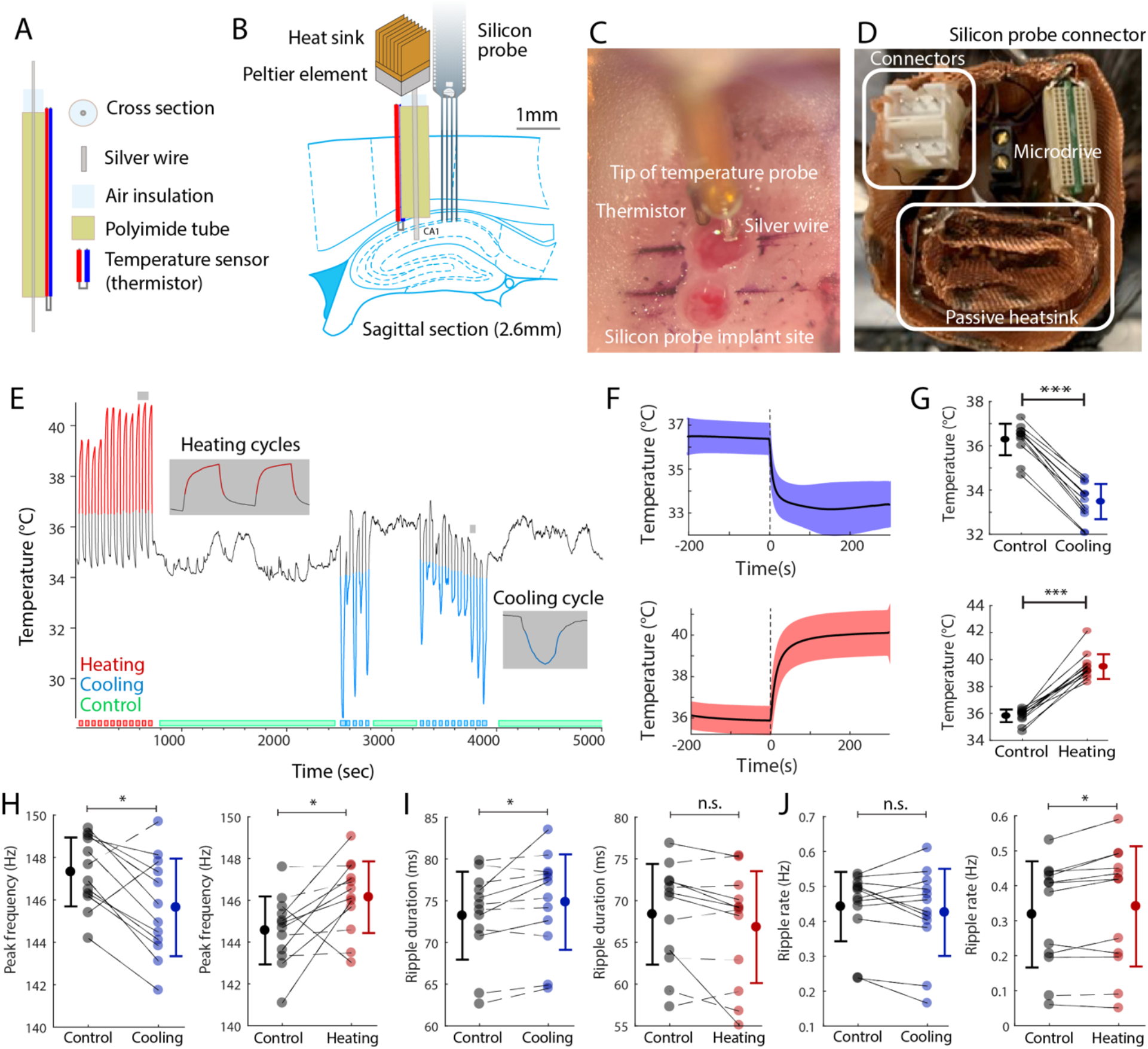
Local temperature manipulation affects ripple frequency. **A-B)** Cooling probe and silicon probe or tungsten wires were implanted in the CA1 region of the hippocampus of rats. Peltier element with heatsink is coupled to the silver wire and the hippocampus is cooled by thermal conduction. **C)** Intraoperative photograph showing the implanted cooling device (top) and the location of the probe implantation (bottom). Black marker lines are approx. 1 mm apart. **D)** Additional copper mesh heatsink is attached to the Peltier element and placed inside the on-head Faraday cage. Custom connectors (for Peltier probe and thermistor) are highlighted on the left. Omnetics connector of the silicon probe and microdrive in black are also shown. **E)** CA1 temperature during local temperature manipulation. Cooling trials are shown by a blue line, and heating trials by red line. Manipulation intervals were defined using the graphical interface StateExplorer (Fig. S1). **F)** Time course of local temperature change during cooling and heating. **G**) Temperature changes during individual cooling and heating sessions (cooling: P = 0.0005, Wilcoxon signed rank test, n = 12 sessions in 3 rats and heating: P = 0.0005, Wilcoxon signed rank test, n = 12 sessions in 4 rats). **H**) Peak frequency of ripples during cooling (left; Δ_freq_ = −1.7Hz, P=0.034) and heating sessions (right; Δ_freq_ = 1.5Hz, P=0.012). Solid lines represent sessions with significant within-session modulation (P < 0.01, Kolmogorov-Smirnov test), and dashed lines represent sessions with non-significant modulation (P > 0.05, Kolmogorov-Smirnov test). **I)** Ripple duration (cooling: Δ_duration_ = 1.6ms, P=0.016; heating: Δ_duration_ = −1.6ms, P=0.077), and **J)** rate of ripple occurrence (cooling: Δ_rate_ = −0.012Hz, P=0.30; heating: Δ_rate_ = 0.02Hz, P=0.027; same sessions shown in G-J; Wilcoxon signed rank test applied in all stats).

## Discussion

We found a correlation between physiological brain temperature variations and the frequency of hippocampal ripples. In addition, we replicated the temperature effect by local cooling and heating the hippocampal CA1 region. The decrease and increase in ripple frequency upon cooling and heating, respectively, suggest that brain temperature is the main mechanism responsible for the ripple frequency differences between sleeping and waking animals. In contrast to changes in ripple frequency, the rate of occurrence of SPW-Rs and ripple duration were only marginally affected by local temperature perturbation, suggesting that the control mechanisms of these parameters reside upstream from the CA1 region.

The effect of ambient temperature on the nervous system and behavior is strong in cold-blooded animals (O’Leary and Marder, 2016). In homoiotherm birds and mammals, while body and brain temperature are homeostatically regulated, there is still a systematic and consistent variation in brain temperature, corresponding to approximately 3°C in birds, rodents and humans (Long and Fee, 2008; Moser and Andersen, 1994, Kiyatkin, 2002) and even much larger changes are present in hibernating animals (Ruf and Geiser, 2015). During sustained nREM sleep, temperature on the neocortical surface of mice decreases by ~2°C (Fuller et al., 1998), and recent work indicates that a specific hypothalamic circuitry exists to deliberately cool the brain and simultaneously induce nREM sleep (Harding et al., 2020, 2021). Our findings confirm previous observations in the rodent regarding the behavior and brain-state dependence of temperature variation (Alföldi et al., 1990; Franken et al., 1992; Hayward and Baker, 1968; Moser et al., 1993).

We extend these previous findings by quantifying the relationship between brain temperature and both SPW-R occurrence, duration, and ripple frequency. The hippocampal SPW-R is a complex pattern of two independent but coupled events. The extracellular sharp wave (SPW) is produced by large transmembrane currents in the apical dendrites of CA1 pyramidal cell, which are triggered by the synchronous CA3 input targeting the mid str. radiatum (Buzsáki et al., 1983). The CA3 volley also excites CA1 interneurons to protract the rate of pyramidal neuron recruitment and their interaction induces a short-lived fast oscillation (110-160 Hz) detected in the local field potential (LFP) as a ‘ripple’ (Buzsáki et al., 1992; O’Keefe and Nadel, 1978; Stark et al., 2014; Ylinen et al., 1995). The two mechanisms can be dissociated by separate perturbations of the CA1 and CA3 regions (Davoudi and Foster, 2019; Nakazawa et al., 2002; Rogers et al., 2021). In our experiments, physiological decrease of brain temperature during nREM sleep was correlated with both the rate of SPW-Rs and ripple frequency, presumably because both the CA1 region and regions upstream to it were cooled. In contrast, artificial manipulation of local CA1 temperature affected ripple frequency but had an inconsistent effect on SPW-R rate. We assume that the minor change of SPW-R rate with temperature increase was due to increasing the temperature also in the CA3 region. This is consistent with previous observations that although the spatial temperature gradient is steep, nevertheless it can have a detectable effect a few millimeters from the probe (Long and Fee, 2008; Petersen and Buzsáki, 2020). The same manipulation of the contralateral CA1 region was without an effect, further supporting the role of local CA1 mechanisms for controlling ripple frequency (Stark et al., 2014). The difference between ripple frequency in the waking and sleeping rat was approximately 5 Hz (Roumis and Frank, 2015). In contrast, perturbation of local CA1 temperature brought about only 2 to 4 Hz shifts. Thus, one may suggest that other factors than cooling play an important role in decreasing ripple frequency during nREM sleep. However, axons of fast firing CA1 basket cells, the presumed substrate of ripple frequency generation (Chiovini et al., 2014; Stark et al., 2014), reach the entire fimbio-subicular extent of CA1 and up to 1 millimeter along the long axis (Sik et al., 1995). Thus, many cell bodies of basket cells residing outside the effectively cooled local patch could have counteracted the frequency decrease brought about by the locally cooled neurons. The primacy of temperature control is further supported by our statistical analyses, which showed that the best single predictor of ripple frequency was temperature, rather than power spectrum slope, brain state, global firing rate or theta/delta ratio of LFP.

How does temperature affect ripple frequency? The key determinant of ripple frequency is the fast-reacting GABA_A_ receptors on the dendrites of basket cells (Stark et al., 2014). Previous work in vitro has demonstrated that cooling the brain slice by only two degrees, the time constant of inhibitory postsynaptic currents in the hippocampus increased by about the same extent as induced by GABAA receptor blockers and general anesthetics at sedative doses (Franks, 2008). Conversely, drugs affecting brain temperature may exert an effect on circuit operations and behavior mediated by direct temperature changes. Auxiliary mechanisms could involve the expression of the cold-inducible RNA-binding protein (CIRBP) and RNA-binding motif protein 3 genes (Morf et al., 2012). However, the proteins encoded by these genes are likely required for structural remodeling and their involvement in the fast communication among interneurons has yet to be uncovered. In summary, the physiological changes of brain temperature between waking and nREM sleep appears to be the major mechanism for altering the frequency of hippocampal ripples.

## Supplementary Material

### Supplementary Figures

**Suppl. Fig. 1:**
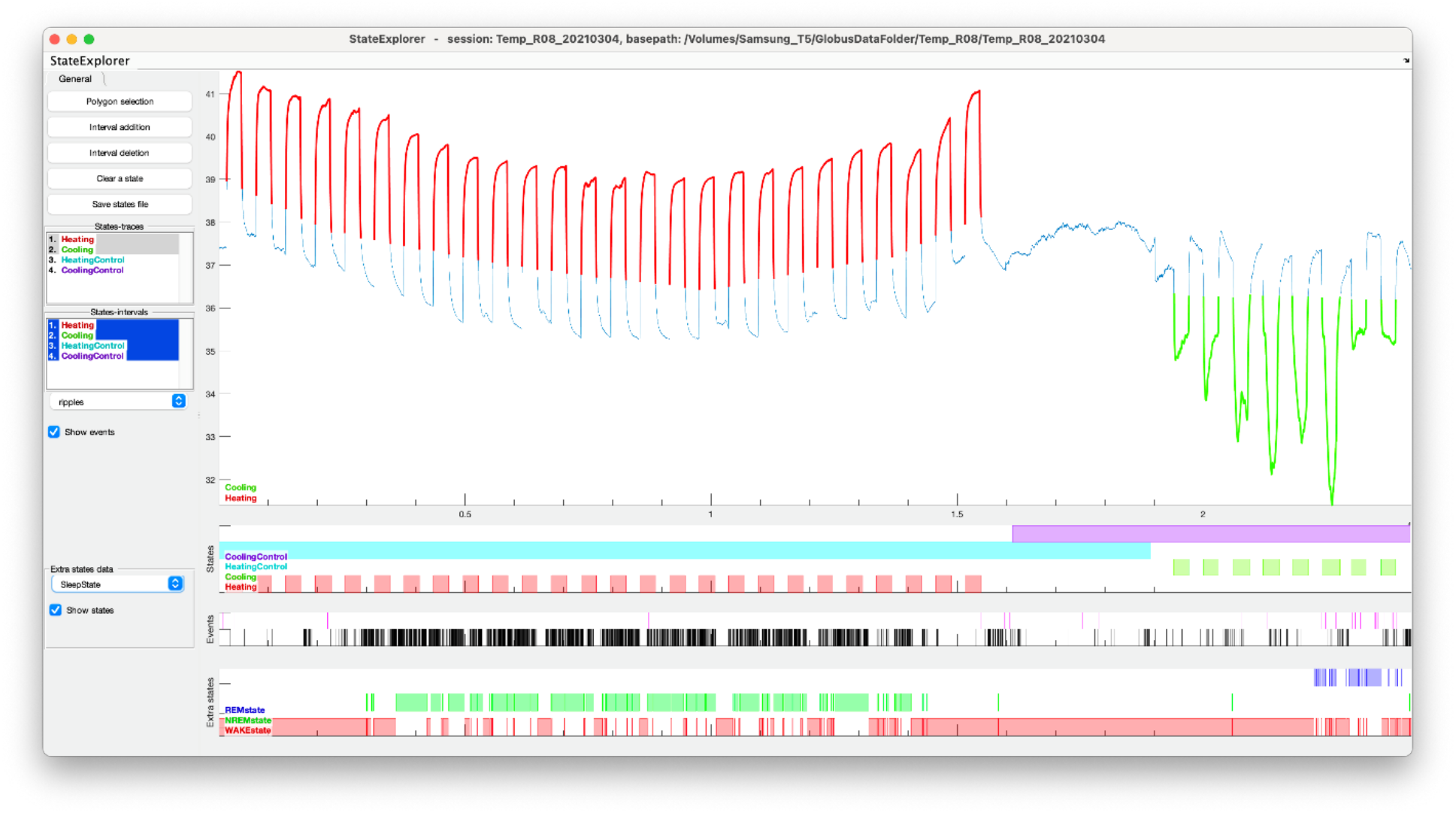
StateExplorer. A graphical user interface to explore, edit and tag brain, behavioral and manipulation states. State scoring on a nonstationary baseline is difficult, we therefore developed a graphical interface based on the data standards of CellExplorer (Petersen et al., 2021), which allows for manual curation of states. Here we characterized the sessions into *Cooling, CoolingControl, Heating* and *HeatingControl*. The GUI is written in Matlab and consists of a left side panel allowing creation of states using temporal borders as well as polygon drawings. The top traces show the brain temperature from a session which consists of both heating and cooling trials. The red lines correspond to heating intervals and the green lines to cooling. The second panel shows the temporal intervals of the same characterized states. Third panel shows ripple events with black rasters. The lowest panel shows the brain states (REM, nonREM, and awake states are respectively shown in blue, green, and red).

**Suppl. Fig. 2:**
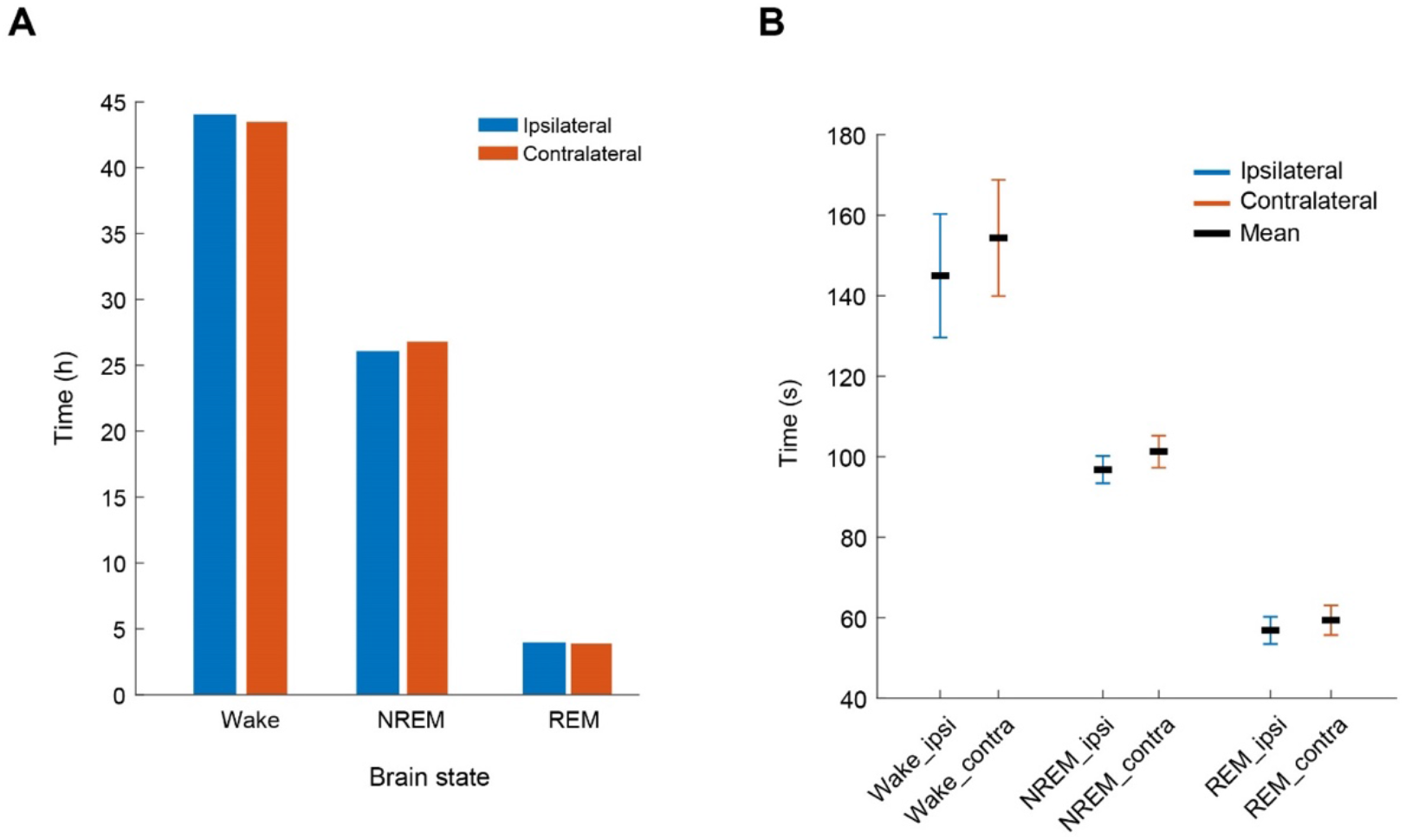
Local temperature manipulation does not affect brain states. Brain state scoring was performed using an ipsilateral and a contralateral recording site. Brain states were categorized as wake, NREM and REM sleep epochs (Watson et al., 2016). **A)** Total time spent in wake, NREM and REM sleep using either the ipsilateral and contralateral brain activity (44.04 and 44.47 hours, 26.06 and 26.78 hours, 3.98 and 3.87 hours; n = 8 sessions in 2 rats). **B)** Average time spent in the 3 different brain states (mean ± SEM).

**Suppl. Fig. 3:**
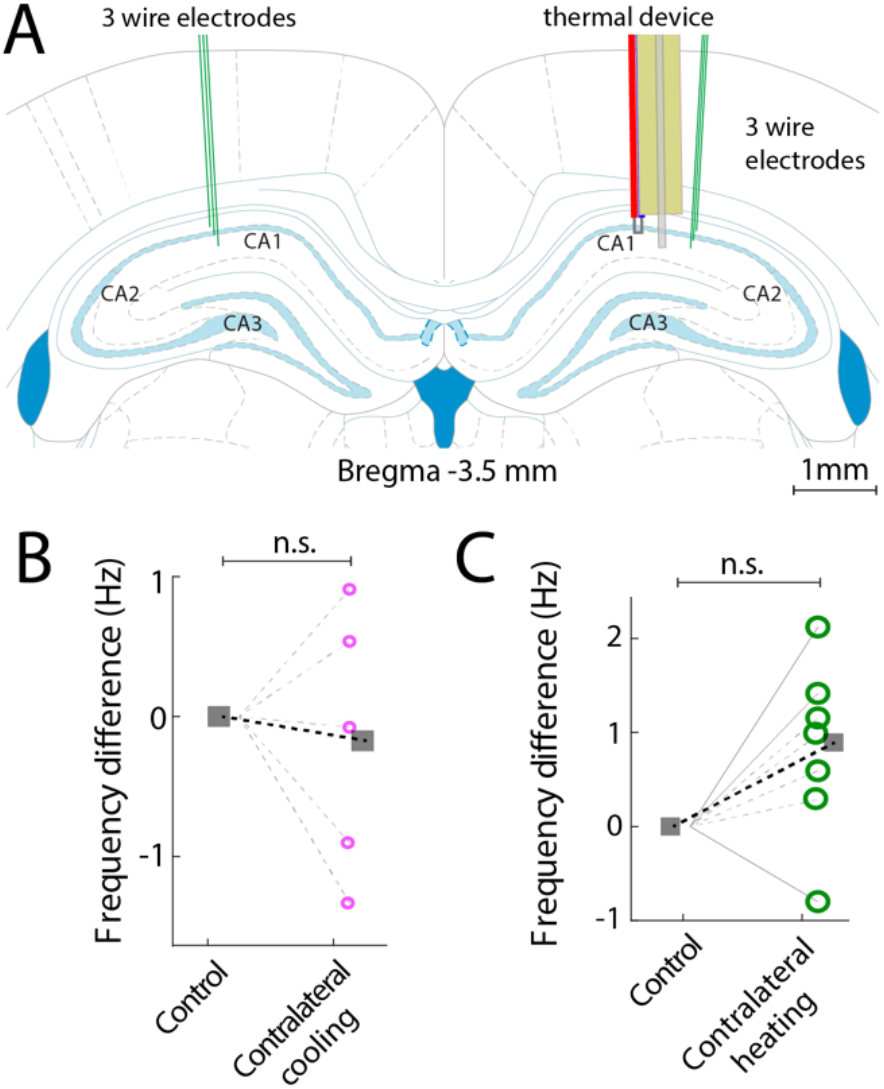
Contralateral temperature perturbation does not affect ripple frequency. **A.** Sketch with bilateral CA1 electrode implants together with a thermal manipulation device on the ipsilateral hemisphere. Cooling **(B)** or heating **(C)** in one hemisphere did not affect the ripple peak frequency in the contralateral hippocampus (Cooling: P=0.82, n session = 5). (Heating: P = n = 8 sessions in 2 rats; Kolmogorov–Smirnov test). Note small frequency variations (0 Hz to max Hz).

**Suppl. Table 1:** Summary of animal subjects with the surgical locations, recording and cooling devices.

| **Animal** | **Recording implant** | **Manipulation implant** | **Behavioral protocol** | **Temperature sensor** |
| --- | --- | --- | --- | --- |
| Temp_R04 | Buzsaki-32 | Passive peltier device | Home cage | Thermistor |
| Temp_R05 | A5X12/16-Buz/Lin-5mm-100-200-160/177 H64LP Package | Passive peltier device | Home cage | Thermistor |
| Temp_R07 | Ipsilateral: tungsten triplet is Bregma-3.8 and ML - 2.5mm, 20-degree angle<br>Contralateral: tungsten triplet is Bregma-3.8 and ML + 2.5mm | Passive peltier device | Home cage | Thermistor |
| Temp_R08 | Ipsilateral: tungsten triplet is Bregma-3.5 and ML - 2.5mm, 10-degree angle<br>Contralateral: tungsten triplet is Bregma-3.5 and ML + 2.5mm | Passive peltier device | Home cage | Thermistor |
| Temp_R09 | Ipsilateral: A5X12/16-Buz/Lin-5mm-100-200-160/177 H64LP Package<br>Contralateral: tungsten triplet is Bregma-3.5 and ML + 2.5mm | Passive peltier device | Home cage | Thermistor |
| MS12 | NeuroNexus Buzsaki 5×12 in CA1 in both hemispheres | Dry ice cryoprobe in MS | Theta maze, wheel box | Thermocouple |
| MS13 | Cambridge Neurotech P1-64ch, poly 2, in CA1 in both hemispheres | Dry ice cryoprobe in MS | Theta maze, wheel box | Thermocouple |
| MS21 | 2 x NeuroNexus Buzsaki64sp, 6 shank, 10 staggered sites, implanted in CA1 bilaterally | Dry ice cryoprobe in MS | Theta maze, linear track, wheel box | Thermocouple |
| MS22 | 2 x NeuroNexus Buzsaki64sp, 6 shank, 10 staggered sites, implanted in CA1 bilaterally | Dry ice cryoprobe in MS | Theta maze, linear track, wheel box | Thermocouple |
| R2W3 | 64-ch polyimide probe, University of Michigan | NA | Home cage | Thermistor |

## Methods

### Subjects and surgery

Rats (adult male Long-Evans, 250-450 g, 3-6 months old) were kept in a vivarium on a 12-hour light/ dark cycle and were housed 2 per cage before surgery and individually after it. All experiments were approved by the Institutional Animal Care and Use Committee at New York University Medical Center.

Animals were anesthetized with isoflurane anesthesia and craniotomies were performed under stereotaxic guidance. A custom designed, 3D printed base (Vöröslakos et al., 2021; Suppl. Fig. 1A) was attached to the skull with meta-bond, serving as a base for the probe implants and protection. A 12 cm by 12 cm sheet of copper mesh (Dexmet Corporation, Wallingford, CT) had been attached to the base with dental cement (Pearson Dental, Sylmar, CA) prior to surgery, from which a protecting cap was formed later (Vöröslakos et al., 2021). Rats (Suppl. Table 1) were implanted with either silicon probes or tungsten wire triplets (50 μm diameter) to record local field potential (LFP) and spikes from the CA1 pyramidal layer. The tip of the cooling device was implanted at AP: −2.5 mm, ML: 2.5 mm, and lowered 2.5 mm below the brain surface, after which it was attached to the skull and base. Silicon probes (NeuroNexus, Ann-Arbor, MI and Cambridge Neurotech, Cambridge, UK) were implanted in the dorsal hippocampus (antero-posterior (AP) − 3.5 mm from Bregma and 2.5 mm from the midline along the medial-lateral axis (ML)). Silicon probes were mounted on custom-made micro-drives to allow their precise vertical movement after implantation (Vöröslakos et al., 2021). Probes were implanted above the target region by attaching the micro-drives to the skull with dental cement. Craniotomies were sealed with sterile wax. Stainless steel screws were implanted above the cerebellum, serving as ground and reference, respectively, for electrophysiological recordings. At the end of electrode and cryoprobe implantation (see below), the copper mesh was folded upwards, connected to the ground screw, and painted with dental cement. The mesh acts as a Faraday cage, shielding the recordings from environmental electric noise and muscle artifacts, provides structural stability and keep debris away from the probe implants. After post-surgery recovery, probes were moved gradually in 50 to 150 μm steps until they reached the CA1 the pyramidal layer. The pyramidal layer of the CA1 region was identified by physiological markers: increased unit activity, strong theta oscillations and phase reversal of the sharp wave ripple oscillations (Mizuseki et al., 2011).

### Peltier cooling device

A passively cooled peltier device was attached to a silver wire that conducted cooling to the CA1 region of the hippocampus. The hot side of a peltier device (00301-9X30-10RU2, TE Technology, Inc., Traverse City, MI) was attached to a copper heatsink (5mm × 5 mm, Enzotech MOS-C10 Forged Copper MOSFET Heatsinks) with heat-conductive adhesive (Arctic Silver Thermal Adhesive, Arctic Silver Inc., Visalia, CA). The heatsink was manually expanded and copper mesh was soldered to it to increase the surface to air ratio (surface area). A 15 mm long silver wire (200μm diameter, 782000, A-M systems, Sequim, WA) was attached to the cold side of the peltier device with heat conductive adhesive. A 5-mm long polyimide tube (EW-95820-05, Cole-Parmer, Vernon Hills, IL) was attached around the silver wire, sealed, and a temperature sensor (223Fu3122-07U015, Semitec USA Corp., Torrance, CA) was attached to the tube. 1 mm of the silver wire was exposed at the tip of the cooling device (Fig. 4a).

### Dry ice cryoprobe

A 15-mm diameter 3D printed container with lid was constructed and a hollow cylinder made from Styrofoam (outer diameter: 18 mm, inner diameter: 10 mm) was inserted into the container. A 20 mm silver wire (127 μm diameter, #781500, A-M systems, Sequim, WA), wrapped with graphene sheet (50 μm thickness), was inserted through the base of the styrofoam container and attached to the inside of the chamber with thermal adhesive (Arctic Silver Thermal Adhesive, ASTA-7G). 10 mm of the wire was protruding from the base of the chamber. The protruding wire was then inserted into an 8 mm long hollow polyimide tube (1 mm diameter), such that 1.5 mm of the silver wire was exposed. The polyimide tube was further sealed with thermal adhesive in both ends to create an air vacuum around the silver wire (Fig. 4a). The air vacuum served as a thermal isolation, to minimize the cooling effects along the wire (Aronov and Fee, 2011; Petersen and Buzsáki, 2020). Finally, a thermocouple (5SC-TT-K-40-72, Omega Engineering Inc., Norwalk, CT) was attached with epoxy adhesive to the cooling implant with the tip of the probe aligned with the protruding silver wire (Fig. 4a). Cooling with dry ice was achieved by placing a small amount of dry ice into the “cooling chamber”, which conducted the cooling to the exposed implanted silver wire.

### Electrophysiological Recordings

Animals were handled daily and accommodated to the experimenter before surgery. After recovery from surgery, the animals were recorded in their home cages and on a set of behavioral mazes. The behavioral sessions typically lasted 40 min, while the total recording time ranged from a couple of hours to a full 24-hour session.

### Sharp wave ripple detection

A single LFP signal was bandpass filtered in the ripple band (80Hz-240Hz), and ripples were detected with a fixed minimum amplitude of 48μV and further fulfilling a duration criterion of 20 ms above 18μV. Ripple events with a duration > 150 ms were excluded to minimize artifacts. The detected events were further manually inspected using NeuroScope2 (Petersen et al., 2021), and noise artifacts and false detected events were removed, typically occurring due to electrical artifacts or scratching artifacts. When possible a reference channel outside the CA1 was also used to filter out false positive ripple events.

### Brain temperature

Brain temperature was measured using either a thermistor or a thermocouple type k and recorded using the Intan system analog amplifier at 20KHz. Temperature data was down-sampled first to 1250Hz and to 1Hz for a subset of the analysis.

### Brain state scoring

Brain state scoring. Brain state scoring was performed as in Watson et al., (2016). In short, spectrograms were constructed with a 1s sliding 10s window fast Fourier transform of 1250Hz data at log-spaced frequencies between 1Hz and 100Hz. Three types of signals were used for score state: broadband LFP, narrowband theta frequency LFP and electromyogram (EMG). For broadband LFP signal, principal component analysis was applied to the Z-transformed (1Hz– 100Hz) spectrogram. The first principal component in all cases was based on power in the low (32Hz) frequencies. Theta dominance was taken to be the ratio of the power at 5Hz-10Hz and 2Hz-16Hz from the spectrogram. EMG was extracted from the intracranially recorded signals by detecting the zero time lag correlation coefficients (r) between 300Hz and 600Hz filtered signals (using a Butterworth filter at 300Hz-600Hz with filter shoulders spanning to 275Hz-625Hz) recorded at all sites. Next all states were inspected and curated manually, and corrections were made when discrepancies between automated scoring and user assessment occurred.

## QUANTIFICATION AND STATISTICAL ANALYSIS

Electrophysiological recordings were conducted using an Intan recording system: RHD2000 interface board with Intan 64 channel preamplifiers sampled at 20 kHz (Intan Technologies, Los Angeles, CA).

### Statistical Analyses

All statistical analyses were performed with MATLAB functions or custom-made scripts. For rank order calculation, the probability of participation and firing rate correlations, the unit of analysis was single cells. Unless otherwise noted, for all tests, non-parametric two-tailed Wilcoxon ranksum (equivalent to Mann-Whitney U-test), Wilcoxon signed-rank or Kruskal-Wallis one-way analysis of variance were used. Due to experimental design constraints, the experimenter was not blind to the manipulation performed during the experiment.

## DATA AND CODE AVAILABILITY

The dataset will be available from our data bank (Petersen et al., 2020) via our website: https://buzsakilab.com/wp/database/

## KEY RESOURCES TABLE

| Resource | Source | Identifier/Location | Description |
| --- | --- | --- | --- |
| **SUBJECTS** |  |  |  |
| Rat: Long-Evans | Charles River | Cat#: Crl:LE 006 | 10 male adults |
| **DATA** |  |  |  |
| Hippocampal data | All authors | https://buzsakilab.com/wp/database/<br>(Petersen et al., 2020) | Public data share with data from our lab |
| **SOFTWARE** |  |  |  |
| CellExplorer, NeuroScope2, and StateExplorer |  | [CellExplorer.org/](https://cellexplorer.org/)<br>(Petersen et al., 2021) | Cell-classification pipeline and GUI built in Matlab |
| MATLAB | MathWorks | https://www.mathworks.com/ |  |
| Buzcode | Buzsaki Lab | https://github.com/buzsakilab/buzcode | Matlab analysis tools |
| FMA Toolbox | Michaël Zugaro | http://fmatoolbox.sourceforge.net/ | Matlab toolbox for Freely Moving Animal (FMA) |
| **OTHER** |  |  |  |
| Silicon probe (5x12, 6x10) | NeuroNexus | https://neuronexus.com/ |  |
| Silicon probe (4x16) | Cambridge Neurotech | https://www.cambridgeneurotech.com/ |  |
| Wire electrodes | California Fine Wire | Tungsten 99.95% (100211), insulated with Heavy Polyimide (HML - Green) |  |
| Intan RHD2000 | Intan technologies | http://intantech.com/RHD2000_evaluation_system.html |  |
| Motive tracking system | Optitrack | http://optitrack.com/ | 6 Flex3 camera system |

## Acknowledgements

This work was supported by the NIH, U19 NS107616 & U19 NS104590, the Independent Research Fund Denmark, & Lundbeckfonden Denmark.

## Author Contributions

Conceptualization: PP, MV, GB

Methodology: MV, PP

Investigation: PP, MV

Formal analysis: PP

Supervision: GB

Writing—original draft: GB, PP, MV

## References

Alföldi, P., Tobler, I., and Borbély, A.A. (1990). Sleep regulation in rats during early development. Am J Physiol 258, R634–644.

Andersen, P., and Moser, E.I. (1995). Brain temperature and hippocampal function. Hippocampus 5, 491–498.

Aronov, D., and Fee, M.S. (2011). Analyzing the dynamics of brain circuits with temperature: design and implementation of a miniature thermoelectric device. J. Neurosci. Methods 197, 32–47.

Buzsáki, G. (1989). Two-stage model of memory trace formation: a role for “noisy” brain states. Neuroscience 31, 551–570.

Buzsáki, G., Lai-Wo S., L., and Vanderwolf, C.H. (1983). Cellular bases of hippocampal EEG in the behaving rat. Brain Research Reviews 6, 139–171.

Buzsáki, G., Horváth, Z., Urioste, R., Hetke, J., and Wise, K. (1992). High-frequency network oscillation in the hippocampus. Science 256, 1025–1027.

Chiovini, B., Turi, G.F., Katona, G., Kaszás, A., Pálfi, D., Maák, P., Szalay, G., Szabó, M.F., Szabó, G., Szadai, Z., et al. (2014). Dendritic Spikes Induce Ripples in Parvalbumin Interneurons during Hippocampal Sharp Waves. Neuron 82, 908–924.

Csernai, M., Borbély, S., Kocsis, K., Burka, D., Fekete, Z., Balogh, V., Káli, S., Emri, Z., and Barthó, P. (2019). Dynamics of sleep oscillations is coupled to brain temperature on multiple scales. The Journal of Physiology 597, 4069–4086.

Davoudi, H., and Foster, D.J. (2019). Acute silencing of hippocampal CA3 reveals a dominant role in place field responses. Nat Neurosci 22, 337–342.

Deboer, T. (1998). Brain temperature dependent changes in the electroencephalogram power spectrum of humans and animals. Journal of Sleep Research 7, 254–262.

Diba, K., and Buzsáki, G. (2007). Forward and reverse hippocampal place-cell sequences during ripples. Nat Neurosci 10, 1241–1242.

Dijk, D.J., and Czeisler, C.A. (1995). Contribution of the circadian pacemaker and the sleep homeostat to sleep propensity, sleep structure, electroencephalographic slow waves, and sleep spindle activity in humans. J Neurosci 15, 3526–3538.

Dupret, D., O’Neill, J., Pleydell-Bouverie, B., and Csicsvari, J. (2010). The reorganization and reactivation of hippocampal maps predict spatial memory performance. Nat Neurosci 13, 995–1002.

Franken, P., Tobler, I., and Borbély, A.A. (1992). Sleep and Waking Have a Major Effect on the 24-Hr Rhythm of Cortical Temperature in the Rat. J Biol Rhythms 7, 341–352.

Franks, N.P. (2008). General anaesthesia: from molecular targets to neuronal pathways of sleep and arousal. Nat Rev Neurosci 9, 370–386.

Franks, N.P., and Wisden, W. (2021). The inescapable drive to sleep: Overlapping mechanisms of sleep and sedation. Science.

Fuller, A., Carter, R.N., and Mitchell, D. (1998). Brain and abdominal temperatures at fatigue in rats exercising in the heat. J Appl Physiol (1985) 84, 877–883.

Gillespie, A.K., Maya, D.A.A., Denovellis, E.L., Liu, D.F., Kastner, D.B., Coulter, M.E., Roumis, D.K., Eden, U.T., and Frank, L.M. (2021). Hippocampal replay reflects specific past experiences rather than a plan for subsequent choice.

Girardeau, G., Benchenane, K., Wiener, S.I., Buzsáki, G., and Zugaro, M.B. (2009). Selective suppression of hippocampal ripples impairs spatial memory. Nat Neurosci 12, 1222–1223.

Harding, E.C., Franks, N.P., and Wisden, W. (2020). Sleep and thermoregulation. Curr Opin Physiol 15, 7–13.

Harding, E.C., Ba, W., Zahir, R., Yu, X., Yustos, R., Hsieh, B., Lignos, L., Vyssotski, A.L., Merkle, F.T., Constandinou, T.G., et al. (2021). Nitric Oxide Synthase Neurons in the Preoptic Hypothalamus Are NREM and REM Sleep-Active and Lower Body Temperature. Frontiers in Neuroscience 15.

Hayward, J., and Baker, M. (1968). Role of cerebral arterial blood in the regulation of brain temperature in the monkey. American Journal of Physiology-Legacy Content 215, 389–403.

Hubbard, J., Gent, T.C., Hoekstra, M.M.B., Emmenegger, Y., Mongrain, V., Landolt, H.-P., Adamantidis, A.R., and Franken, P. (2020). Rapid fast-delta decay following prolonged wakefulness marks a phase of wake-inertia in NREM sleep. Nat Commun 11, 3130.

Joo, H.R., and Frank, L.M. (2018). The hippocampal sharp wave-ripple in memory retrieval for immediate use and consolidation. Nat Rev Neurosci 19, 744–757.

Long, M.A., and Fee, M.S. (2008). Using temperature to analyse temporal dynamics in the songbird motor pathway. Nature 456, 189–194.

Maguire, E.A., and Hassabis, D. (2011). Role of the hippocampus in imagination and future thinking. PNAS 108, E39–E39.

McGinty, D., and Szymusiak, R. (1990). Keeping cool: a hypothesis about the mechanisms and functions of slow-wave sleep. Trends in Neurosciences 13, 480–487.

Miyawaki, H., and Diba, K. (2016). Regulation of Hippocampal Firing by Network Oscillations during Sleep. Curr Biol 26, 893–902.

Mizuseki, K., Diba, K., Pastalkova, E., and Buzsáki, G. (2011). Hippocampal CA1 pyramidal cells form functionally distinct sublayers. Nature Neuroscience 14, 1174–1181.

Morf, J., Rey, G., Schneider, K., Stratmann, M., Fujita, J., Naef, F., and Schibler, U. (2012). Cold-Inducible RNA-Binding Protein Modulates Circadian Gene Expression Posttranscriptionally. Science.

Moser, E.I., and Andersen, P. (1994). Conserved spatial learning in cooled rats in spite of slowing of dentate field potentials. J. Neurosci. 14, 4458–4466.

Moser, E., Mathiesen, lacob, and Andersen, P. (1993). Association Between Brain Temperature and Dentate Field Potentials in Exploring and Swimming Rats. Science.

Nádasdy, Z., Hirase, H., Czurkó, A., Csicsvari, J., and Buzsáki, G. (1999). Replay and Time Compression of Recurring Spike Sequences in the Hippocampus. J. Neurosci. 19, 9497–9507.

Nakazawa, K., Quirk, M.C., Chitwood, R.A., Watanabe, M., Yeckel, M.F., Sun, L.D., Kato, A., Carr, C.A., Johnston, D., Wilson, M.A., et al. (2002). Requirement for hippocampal CA3 NMDA receptors in associative memory recall. Science 297, 211–218.

Norimoto, H., Makino, K., Gao, M., Shikano, Y., Okamoto, K., Ishikawa, T., Sasaki, T., Hioki, H., Fujisawa, S., and Ikegaya, Y. (2018). Hippocampal ripples down-regulate synapses. Science.

O’Keefe, J., and Nadel, L. (1978). The Hippocampus as a Cognitive Map (Oxford: Clarendon Press).

Olafsdóttir, H.F., Bush, D., and Barry, C. (2018). The Role of Hippocampal Replay in Memory and Planning. Curr Biol 28, R37–R50.

O’Leary, T., and Marder, E. (2016). Temperature-Robust Neural Function from Activity-Dependent Ion Channel Regulation. Curr Biol 26, 2935–2941.

Petersen, P.C., and Buzsáki, G. (2020). Cooling of Medial Septum Reveals Theta Phase Lag Coordination of Hippocampal Cell Assemblies. Neuron 107, 731–744.e3.

Petersen, P.C., Hernandez, M., and Buzsáki, G. (2020). The Buzsaki Lab Databank - Public electrophysiological datasets from awake animals (Zenodo).

Petersen, P.C., Siegle, J.H., Steinmetz, N.A., Mahallati, S., and Buzsáki, G. (2021). CellExplorer: A framework for visualizing and characterizing single neurons. Neuron.

Pfeiffer, B.E., and Foster, D.J. (2013). Hippocampal place-cell sequences depict future paths to remembered goals. Nature 497, 74–79.

Rogers, S., Rozman, P.A., Valero, M., Doyle, W.K., and Buzsáki, G. (2021). Mechanisms and plasticity of chemogenically induced interneuronal suppression of principal cells. PNAS 118.

Roumis, D.K., and Frank, L.M. (2015). Hippocampal sharp-wave ripples in waking and sleeping states. Curr Opin Neurobiol 35, 6–12.

Roux, L., Hu, B., Eichler, R., Stark, E., and Buzsáki, G. (2017). Sharp wave ripples during learning stabilize the hippocampal spatial map. Nat Neurosci 20, 845–853.

Ruf, T., and Geiser, F. (2015). Daily torpor and hibernation in birds and mammals. Biol Rev Camb Philos Soc 90, 891–926.

Schacter, D.L., Addis, D.R., and Buckner, R.L. (2007). Remembering the past to imagine the future: the prospective brain. Nat Rev Neurosci 8, 657–661.

Schomburg, E.W., Anastassiou, C.A., Buzsáki, G., and Koch, C. (2012). The Spiking Component of Oscillatory Extracellular Potentials in the Rat Hippocampus. J Neurosci 32, 11798–11811.

Sheroziya, M., and Timofeev, I. (2015). Moderate Cortical Cooling Eliminates Thalamocortical Silent States during Slow Oscillation. J. Neurosci. 35, 13006–13019.

Sik, A., Penttonen, M., Ylinen, A., and Buzsaki, G. (1995). Hippocampal CA1 interneurons: an in vivo intracellular labeling study. J. Neurosci. 15, 6651–6665.

Stark, E., Roux, L., Eichler, R., Senzai, Y., Royer, S., and Buzsáki, G. (2014). Pyramidal cell-interneuron interactions underlie hippocampal ripple oscillations. Neuron 83, 467–480.

Tingley, D., McClain, K., Kaya, E., Carpenter, J., and Buzsáki, G. (2021). A metabolic function of the hippocampal sharp wave-ripple. Nature 597, 82–86.

Vanderwolf, C.H. (1969). Hippocampal electrical activity and voluntary movement in the rat. Electroencephalography and Clinical Neurophysiology 26, 407–418.

Vöröslakos, M., Miyawaki, H., Royer, S., Diba, K., Yoon, E., Petersen, P.C., and Buzsáki, G. (2021). 3D-printed Recoverable Microdrive and Base Plate System for Rodent Electrophysiology. Bio-Protocol 11, e4137–e4137.

Watson, B.O., Levenstein, D., Greene, J.P., Gelinas, J.N., and Buzsáki, G. (2016). Network homeostasis and state dynamics of neocortical sleep. Neuron 90, 839–852.

Wilson, M.A., and McNaughton, B.L. (1994). Reactivation of Hippocampal Ensemble Memories During Sleep. Science.

Ylinen, A., Soltész, I., Bragin, A., Penttonen, M., Sik, A., and Buzsáki, G. (1995). Intracellular correlates of hippocampal theta rhythm in identified pyramidal cells, granule cells, and basket cells. Hippocampus 5, 78–90.

